# Robust normalization and transformation techniques for constructing gene coexpression networks from RNA-seq data

**DOI:** 10.1101/2020.09.22.308577

**Authors:** Kayla A Johnson, Arjun Krishnan

## Abstract

**Background:** Constructing gene coexpression networks is a powerful approach for analyzing high-throughput gene expression data towards module identification, gene function prediction, and disease-gene prioritization. While optimal workflows for constructing coexpression networks – including good choices for data pre-processing, normalization, and network transformation – have been developed for microarray-based expression data, such well-tested choices do not exist for RNA-seq data. Almost all studies that compare data processing/normalization methods for RNA-seq focus on the end goal of determining differential gene expression.

**Results:** Here, we present a comprehensive benchmarking and analysis of 30 different workflows, each with a unique set of normalization and network transformation methods, for constructing coexpression networks from RNA-seq datasets. We tested these workflows on both large, homogenous datasets (Genotype-Tissue Expression project) and small, heterogeneous datasets from various labs (submitted to the Sequence Read Archive). We analyzed the workflows in terms of aggregate performance, individual method choices, and the impact of multiple dataset experimental factors. Our results demonstrate that between-sample normalization has the biggest impact, with trimmed mean of M-values or upper quartile normalization producing networks that most accurately recapitulate known tissue-naive and tissue-specific gene functional relationships.

**Conclusions:** Based on this work, we provide concrete recommendations on robust procedures for building an accurate coexpression network from an RNA-seq dataset. In addition, researchers can examine all the results in great detail at https://krishnanlab.github.io/norm_for_RNAseq_coexp to make appropriate choices for coexpression analysis based on the experimental factors of their RNA-seq dataset.

## Background

Constructing gene coexpression networks is a powerful and widely-used approach for analyzing high-throughput gene expression data from microarray and RNA-seq technologies [1]. Coexpression networks provide a framework for summarizing multiple transcriptomes of a particular species, tissue, or condition as a graph where each node is a gene and each edge between a pair of genes represents the similarity of their patterns of expression. Coexpressed genes are highly likely to be transcriptionally co-regulated and are often functionally related to each other by virtue of taking part in the same biological process or physiological trait [2]. Many studies have leveraged these properties to use coexpression networks in several important applications such as determining co-regulated gene groups [3] and associating genes to functions and phenotypes [1].

Best practices for normalization when building a coexpression network from a raw gene-expression dataset have been developed and compared for data from microarrays [4,5]. However, coexpression network analysis is now being routinely applied to the exponentially increasing number of data from RNA-seq even though the optimal procedure for network building has not been evaluated and honed for RNA-seq data, particularly in regard to normalization and transformation. Although many normalization strategies have been developed for RNA-seq data, they have mostly been benchmarked only in the context of estimating differential gene expression [6–10]. Very little work has been done so far to comprehensively compare these strategies for normalization and transformation (and their combinations) to construct the most accurate coexpression networks from RNA-seq data, especially to ensure their robust application to datasets typically generated by individual research groups [1].

The most relevant prior work focuses on establishing best practices that reduce the introduction of artifacts in coexpression networks built from RNA-seq data [11]. This study includes a sequential comparison of a select number of methods for transcript assembly, normalization, and network reconstruction. However, the normalization comparison is based on 10 RNA-seq datasets, leaving considerable room for improvement. First is to increase the number and diversity of datasets studied. This is vital for finding robust procedures that work across datasets that can vary considerably in many respects, including sample size, sample variability, sequencing depth, tissue type, and other experimental factors. Further, testing on a wide range of datasets is critical both for the analysis of individual datasets as well as integrative analysis of hundreds/thousands of datasets. Second, not only do more normalization and network transformation methods need to be compared but how they might interact in combinations needs be studied. Third, the resulting networks need to be evaluated directly on the accuracy of the coexpression between gene pairs, instead of performance in a downstream task such as gene function prediction, to ensure maximal utility of the network regardless of the subsequent biological application. Finally, the evaluation metric needs to be informative considering the fact that only a small fraction of all gene pairs in the genome are functionally related and that many of these relationships occur in a tissue-specific manner.

In this work, we present the most comprehensive benchmarking of commonly used within- and between-sample normalization strategies and network transformation methods for constructing accurate coexpression networks from human RNA-seq data. We tested every possible combination of methods from different normalization and transformation stages. Our primary interest is in identifying robust combinations of methods that consistently result in coexpression networks that accurately capture general and tissue-specific gene relationships across a large variety of datasets. This will allow us to propose general recommendations useful for experimental research groups analyzing their own RNA-seq data as well as computational researchers seeking to build many coexpression networks from publicly available data for the purposes of data/network integration. Towards this aim, we use hundreds of datasets, generated by a consortium and by individual laboratories, covering multiple experimental factors. We then test the resulting networks on both tissue-naive and tissue-specific prior knowledge about gene functional relationships. Based on these extensive analyses, we finally provide concrete recommendations for normalization and transformation choices in RNA-seq coexpression analysis.

## Results

### Expression data, gold standard, and benchmarking summary

To test various within-sample normalization, between-sample normalization, and network transformation methods (and their combinations) on a large data collection, we started with gene count data from the Recount2 database [12]. Recount2 contains data from both the Genotype-Tissue Expression (GTEx) project [13] and the Sequence Read Archive (SRA) [14] repository that have been uniformly quality-controlled, aligned, and quantified to the number of reads per gene in the genome. Datasets from the GTEx project allowed us to assess method performance on large, relatively homogeneous datasets with high-sequencing depth and quality. The GTEx data was also critical for investigating the impact of experimental factors such as sample size, which we performed by doing multiple rounds of random sampling from GTEx datasets. Datasets from SRA, on the other hand, were representative of heterogeneous, mostly small experiments (median of 12 samples) that are generated by individual labs, with a range of sequencing depths and qualities. In total, we used 9,657 GTEx samples and 6,301 SRA samples from a total of 287 datasets (**Table 1, Fig. S1**; see *Methods*), and processed and evaluated these two collections separately.

**Table 1:** Summary of data used in this study. See *Figure S1* and *Methods* for more details.

|  | GTEx | SRA |
| --- | --- | --- |
| Data size | 9,657 samples | 6,301 samples / 256 datasets |
| Number of tissues | 31 tissues/datasets | 19 tissues |
| Median dataset size | 197 samples | 12 samples |
|  | Total: 15,958 samples from 37 unique tissues |  |

After preprocessing each dataset using lenient filters in order to keep data for as many genes and samples as possible (see *Methods*), we compared methods commonly used in RNA-seq analysis to effectively construct one coexpression network per dataset (i.e. building 31 GTEx networks and 256 SRA networks). We focused on three key stages of data processing and network building: a) within-sample normalization: counts per million (CPM), transcripts per million (TPM), and reads per kilobase per million (RPKM), b) between-sample normalization: quantile (QNT), trimmed mean of M-values (TMM), and upper quartile (UQ), and c) network transformation: weighted topological overlap (WTO) and context likelihood of relatedness (CLR). To systematically examine these methods and their interactions, we built 30 different workflows covering all possible combinations of choices (**Fig. 1**). For clarity, in the rest of the manuscript, we present individual methods in regular font (e.g. TPM normalization) and italicize workflows (e.g. *TPM*, which is TPM combined with no between-sample normalization and no network transformation, or *TPM_CLR*, which is TPM paired with just CLR).

**Figure 1.**
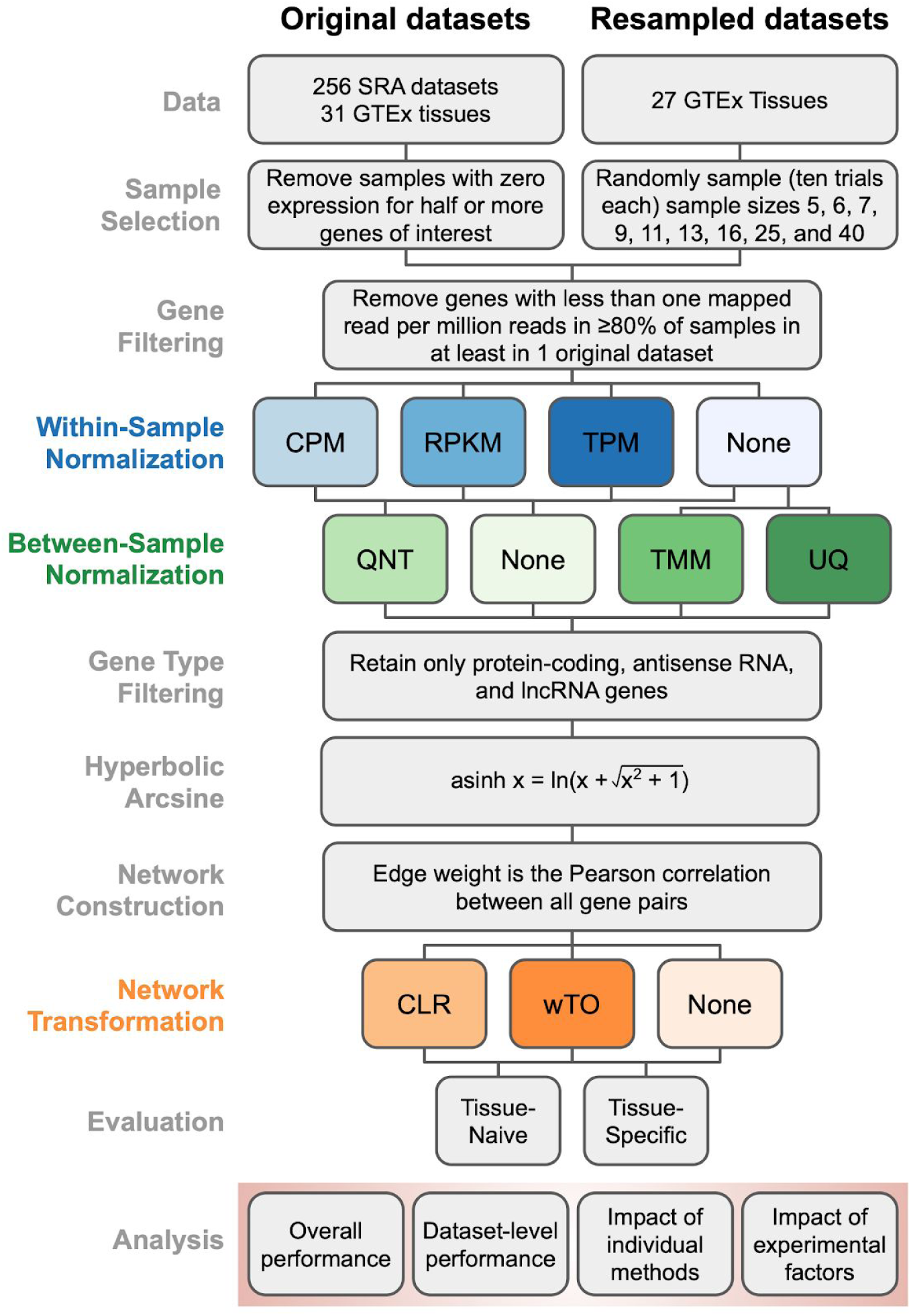
Pipeline for benchmarking the optimal workflow for constructing coexpression networks from RNA-seq data. The main pipeline was executed for the original GTEx and SRA datasets and a large collection of datasets of different sizes resampled from the GTEx datasets. Three key stages within-sample normalization, between-sample normalization, and network transformation – where we tested method choices are highlighted in different colors. All the other stages were composed of standard selection, filtering, and data transformation operations. The coexpression networks resulting from all the workflows were evaluated using two gold-standards that capture generic (tissue-naive) and tissue-specific gene functional relationships. Finally, all the evaluation results were used to analyze the impact of various aspects of the workflows, methods, and datasets on the accuracy of coexpression networks.Abbreviations: CPM (Counts Per Million), RPKM (Reads Per Kilobase Million), TPM (Transcripts Per Million), QNT (quantile), TMM (Trimmed Mean of M-values), UQ (Upper Quartile), CLR (Context Likelihood of Relatedness), WTO (Weighted Topological Overlap).

Since this entire workflow is unsupervised, i.e. not reliant on prior knowledge about gene relationships, we evaluated the resulting coexpression networks by comparing them to gold standards of known gene functional relationships. The gold standards were built using experimentally-verified co-annotations to specific biological process terms in the Gene Ontology [15]. These comparisons yielded evaluation metrics that summarize how well the patterns of coexpression captured in the network reflect known gene functional relationships (see *Network Evaluation* in *Methods*). Further, gene activities and interactions vary drastically depending on cell type or tissue. Hence, we also created tissue-specific gold standards to assess whether the resulting networks were able to recapitulate tissue-specific coexpression in addition to general “tissue-naive” coexpression. Tissue-specific gold standards were created for as many tissues as possible by subsetting the naive gold standard using genes known to be expressed in a particular tissue. While area under the receiver operator curve (auROC) is frequently used to estimate network accuracy, it does not account for the fact that only a small fraction of gene pairs (out of the total possible) biologically interact. In the gold standard, this imbalance is reflected by the number of negatives (non-interactions) far outnumbering the positives (interactions) [16]. Therefore, we measured network accuracy using area under the precision recall curve (auPRC), which emphasizes the accuracy of top-ranked coexpression gene pairs [17].

In total, for each of the 287 datasets from GTEx and SRA, we built one coexpression network per dataset using each of the 30 workflows, resulting in 8,610 coexpression networks. Later on, we create 2,430 additional datasets generated by resampling GTEx that, which when run through all the workflows, resulted in another 72,900 networks. Each GTEx network contains 20,418 genes while each SRA network contains 22,084 genes, and all networks are fully connected with edges weighted by their strength of correlation. Each of these networks were evaluated using the tissue-naive gold-standard and, whenever applicable, the tissue-specific gold-standard.

### Overall performance of workflows

For all 30 workflows, **Figure 2** shows the overall performance of the networks resulting from GTEx (left) and SRA (right) datasets based on evaluation using the tissue-naive gold standard. **Figure S2** shows the performance of these networks based on the tissue-specific gold standards (when available). Overall, networks built from GTEx datasets are far more accurate than those built from SRA datasets (**Fig. 2, S2**). In each of the four cases – GTEx and SRA networks evaluated using tissue-naive and tissue-specific gold standards – the top-performing workflows always contain TMM or UQ normalization. Further transforming the network with CLR (*TMM_CLR* and *UQ_CLR*) results in top-tier workflows for the GTEx datasets regardless of gold standard. However, CLR transformation is only among top-performing methods for SRA datasets in recovering tissue-specific gene relationships. Though *TMM_CLR* and *UQ_CLR* still perform reasonably well on the tissue-naive standard for SRA, there is a clear gap from the top tier. Despite the other between-sample normalization methods overall resulting in good performance, workflows that include quantile normalization (QNT) are conspicuously absent among the top ten workflows for both GTEx and SRA.

**Figure 2.**
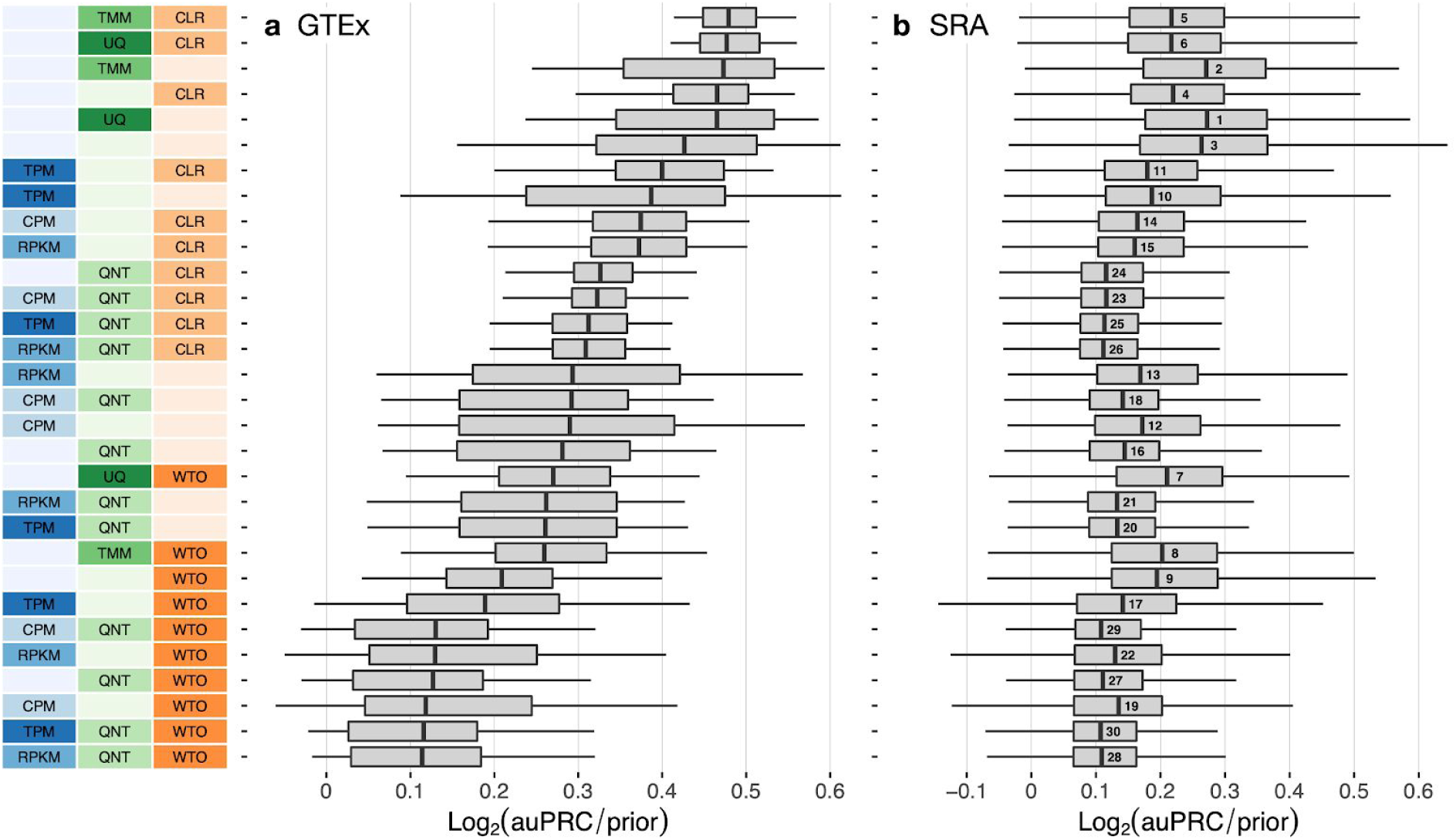
Overall performance of workflows. The plots show the aggregate accuracy of all coexpression networks resulting from each individual workflow using (**a**) GTEx and (**b**) SRA datasets, evaluated based on the tissue-naive gold standard. The workflows (rows) are described in terms of the specific method used in the within-sample normalization (blues), between-sample normalization (greens), and network transformation (oranges) stages. The performance of each workflow is presented as boxplots that summarizes the log2(auPRC/prior) of each workflow where auPRC is the area under the precision recall curve (see *Methods*). The workflows are ordered by their median log2(auPRC/prior) for the GTEx data. The numbers inside the SRA boxes indicate rank by median log2(auPRC/prior) of the workflows for the SRA data. *Figure S2* contains these plots based on the tissue-specific gold standard.

The next noteworthy observation is that the top workflows do not include a within-sample normalization step. Yet, workflows that do include within-sample normalization methods (CPM, RPKM, TPM) can perform better than many other workflows depending on other choices made in the pipeline, the best choice often is to be paired with no other method or CLR alone. For GTEx datasets, CLR seems to either have no effect or result in slightly improved performance, while the WTO transformation almost exclusively makes up the bottom tier of workflows. For building networks from SRA datasets, although workflows including WTO do not exclusively end up in the bottom tier (as is the case with GTEx data), adding WTO to a particular workflow always hurts performance. The worst workflows for SRA in either standard are quantile normalization (QNT) paired with CLR or WTO.

### Dataset-level performance of workflows

Next, we dissected the aggregated results described above for GTEx and SRA as a whole by examining the accuracy of these workflows on a per-dataset basis. First, we compared pairs of workflows to each other and determined the proportion of datasets in which one workflow outperformed the other across all GTEx and all SRA datasets (**Fig. 3, S3–5** heatmap colors). Second, we performed paired statistical tests to estimate the significance of the difference between the workflows (**Fig. 3, S3–5**, asterisks on the heatmap). Finally, we scored each workflow based on the number of other workflows it significantly outperforms (**Fig. 3, S4** barplots). Based on this analysis, in the ‘GTEx-naive’ setting (i.e. networks from GTEx data evaluated on the tissue-naive gold standard), we observed that four workflows are all significantly more accurate than 25 other workflows but not significantly different from one another (paired Wilcoxon rank-sum test; corrected p-value < 0.01; **Fig. 3**). Within these four workflows, *TMM* outperforms *TMM_CLR, UQ*, and *UQ_CLR* on 58%, 61%, and 58% of GTEx networks, respectively. The *TMM* workflow is also significantly better most number of times compared to other workflows in the SRA networks using the naive standard, although *Counts* and *UQ* are only slightly behind *TMM* (**Fig. 3, S3**). These workflows tie for first place when SRA networks are evaluated on the tissue-specific gold standards (**Fig. S4, S5**).

**Figure 3.**
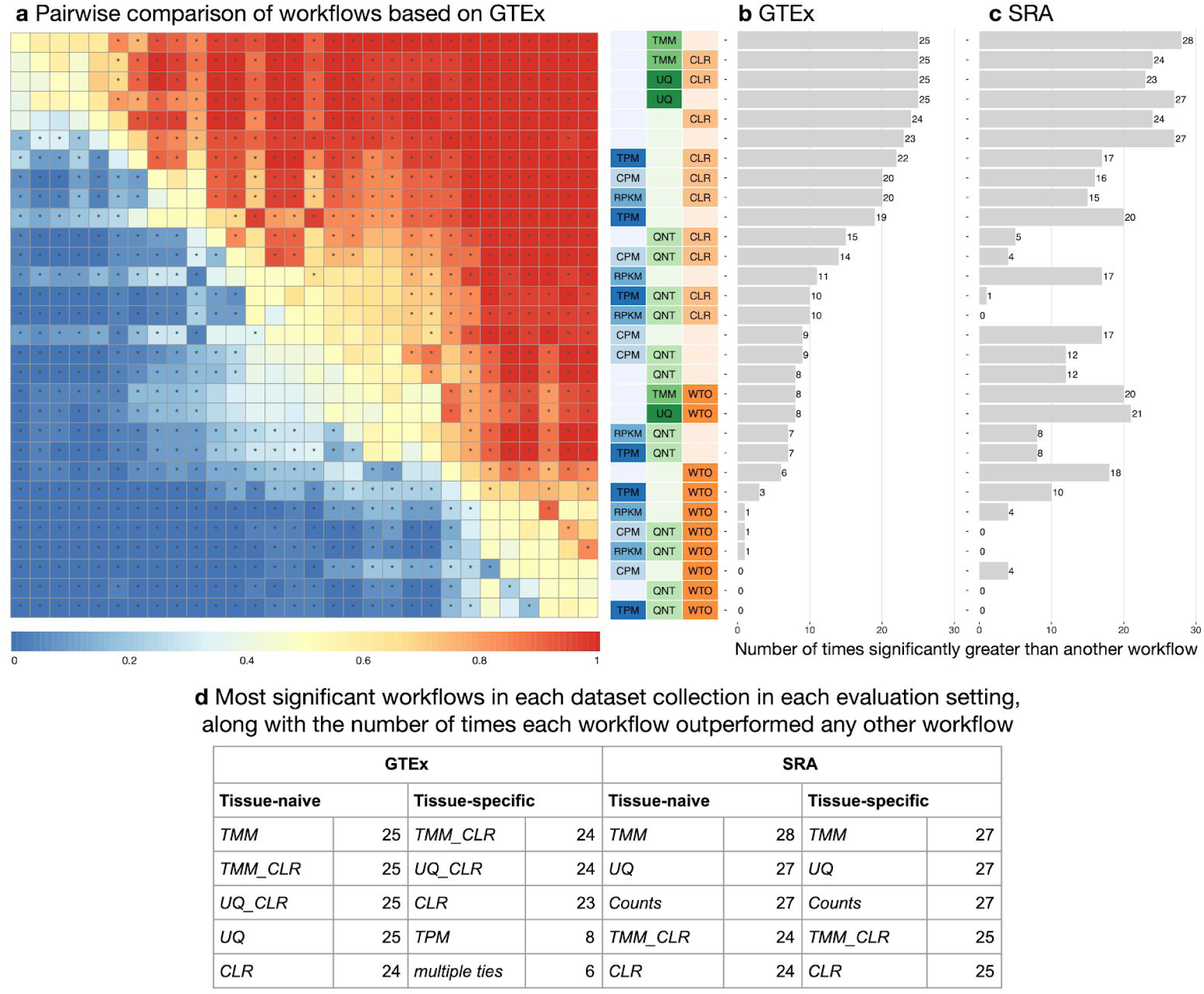
Dataset-level pairwise comparison of workflow performance. (**a**) The heatmap shows the relative performance of a pair of workflows, corresponding to a row and a column, directly compared to each other for the GTEx datasets based on the tissue-naive gold standard. The workflows along the rows are depicted using color swatches similar to *Figure 2*. The color in each cell (row, column) represents the proportion of datasets for which the workflow along the row has a higher log2(auPRC/prior) than the workflow along the column. Comparisons that are statistically significant (corrected p < 0.01) based on a paired Wilcoxon test are marked with an asterisk. *Figure S3* contains the corresponding heatmap for the SRA datasets. (**b** and **c**) Barplots show the number of times each workflow was significantly greater than another workflow for GTEx and SRA datasets. *Figures S4* and *S5* contain these performance plots based on the tissue-specific gold standard. (**d**) The table shows the most significant workflows across evaluation cases along with the number of times a given workflow outperformed any other workflow for the GTEx and SRA datasets based on the tissue-naive and tissue-specific gold standards.

When the GTEx networks are evaluated on tissue-specific standards, there are much fewer significant differences between workflows overall, with the exception of *TMM_CLR, UQ_CLR*, and *CLR* being significantly greater than 23 or 24 workflows (**Fig. S4**). Here, *TMM_CLR* performs better than *UQ_CLR* on 57% of networks and better than *CLR* on 76% of networks. Despite having similar median log2(auPRC/prior) values to *TMM_CLR* and *UQ_CLR* (**Fig. S2**), the *UQ* and *TMM* workflows only perform significantly better than another workflow a handful of times (**Fig. S4**). This suggests that including CLR in the workflow is especially helpful in capturing tissue-specific coexpression in the GTEx networks.

Again, the impact of within-sample normalization varies depending on the choice of the other methods in the workflow. *TPM_CLR* is generally the top-performing workflow among those including within-sample normalization across evaluation cases, though *TPM* slightly outperforms *TPM_CLR* for the SRA networks evaluated on the naive standard (**Fig. 3** and **S3**).

The impact of network transformation is similar between GTEx and SRA data, but there is disagreement in the very top method. With GTEx, workflows that include CLR tend to be significant the most number of times, while WTO-containing workflows tend to be the least. Not a single workflow with WTO significantly outperformed any other workflow for GTEx based on the tissue-specific gold standard (**Fig. S4**). On the other hand, CLR workflows perform well on the SRA networks, but do not constitute the workflows that were significantly greater than another the absolute most number of times (**Fig. S3** and **S5**). WTO hurts performance in every case even here. Pairing either CLR or WTO with quantile normalization (QNT) yields particularly poor performance in the SRA networks. All together, these results suggest that TMM yields the most accurate coexpression network by a very close margin and CLR can further improve the network in select cases.

### Impact of individual methods on performance of workflows

Though the previous analyses shed light on the contributions of individual methods, we wanted to more explicitly assess how choosing or not choosing a particular within-sample normalization, between-sample normalization, or network transformation affects general performance of any given workflow. To this end, for each method, we calculated the proportion of times that workflows that include a particular method performed significantly better than workflows that did not include the method (**Fig. 4**; see *Methods* for details).

**Figure 4.**
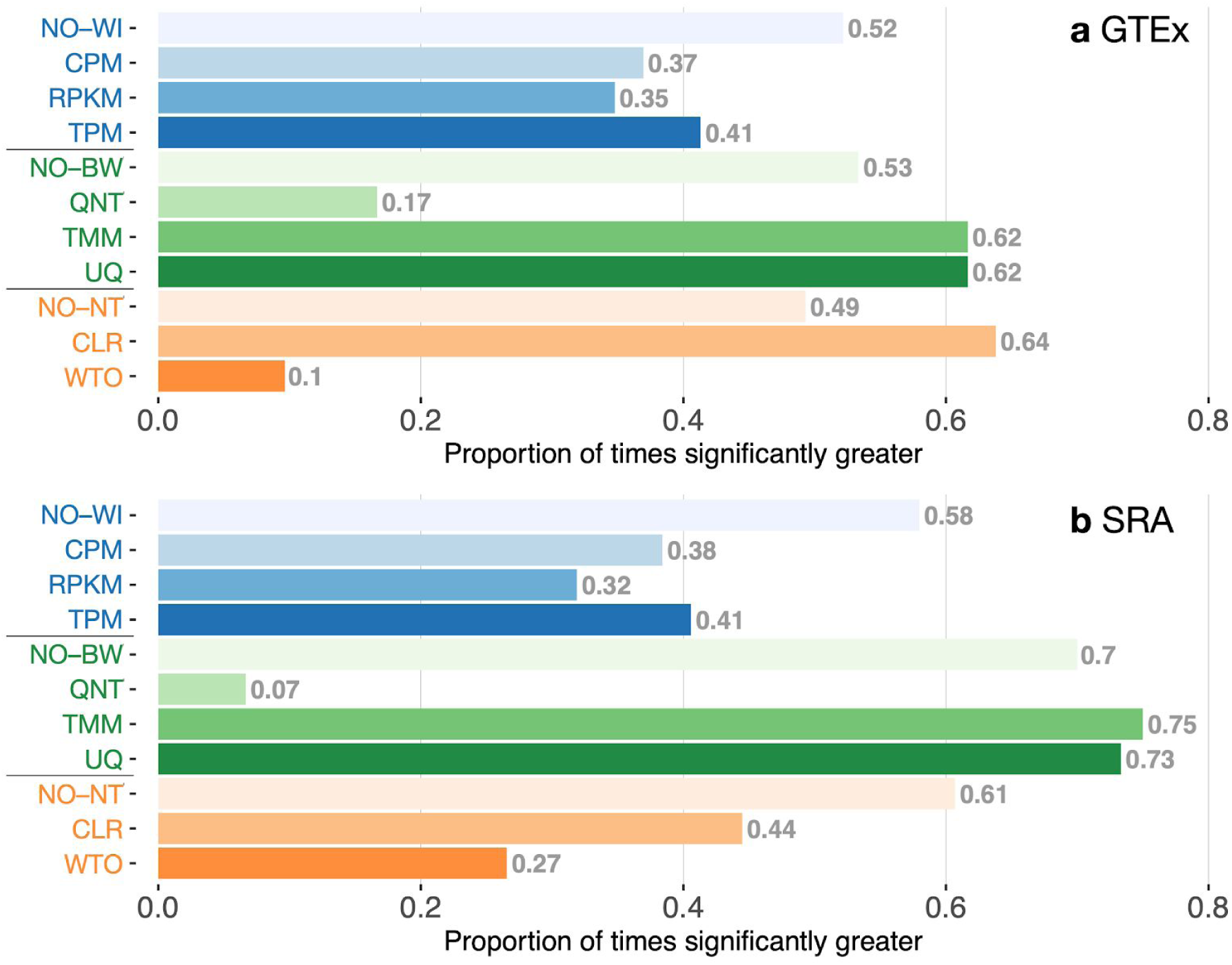
Impact of individual methods on performance of workflows. Each bar in the two barplots, corresponding to a specific method, shows the proportion of times (x-axis) that workflows including that particular method (y-axis) were significantly better than other workflows. The barplots correspond to performance for the (**a**) GTEx and (**b**) SRA datasets evaluated on the tissue-naive gold standard. In order to make the comparison of between-sample normalization methods fair, workflows also including CPM, RPKM, or TPM were left out because it is not possible to pair them with TMM or UQ normalization. Similarly, TMM and UQ methods are not included for “no within-sample normalization” (NO–WI). *Figure S6* contains these barplots based on the tissue-specific gold standard.

This analysis clearly shows that, in all four cases (GTEx and SRA, each with tissue-naive and tissue-specific standards), utilizing any within-sample normalization method results in worse overall performance than not using it (**Fig. 4** and **S6**). Among within-sample normalization methods, TPM usually performs slightly better than CPM and RPKM. TMM and UQ are the best between-sample normalization methods. Their performances are exactly equal for GTEx data evaluated on either standard, and TMM is slightly better than UQ for SRA data in both evaluations. However, doing no between-sample normalization performs quite well too, only narrowly worse than TMM or UQ. It is clear in all four cases that quantile normalization (QNT) is vastly outperformed. Network transformation is the group most obviously different between GTEx and SRA data, with CLR being the clear winner for GTEx, while not doing any network transformation is significant many more times for SRA regardless of gold standard (**Fig. 4 and S6**).

### Impact of varying experimental factors on performance of workflows

The reason we included SRA data in this study is that SRA datasets are very representative of expression datasets typically generated by numerous individual laboratories. Accordingly, these datasets vary considerably in terms of multiple factors including sample size, sample similarity, number of mapped reads, and tissue type. Though these factors impact the quality of coexpression networks derived from the individual datasets, it is hard to tease out the effect of each of these factors (controlling for others) on the accuracies that we observed using different workflows on SRA data. Therefore, using the large GTEx datasets, we created a collection of SRA-like datasets to more closely examine the impact of each experimental factor. First, we determined the nine sample sizes (5, 6, 7, 9, 11, 13, 16, 25, and 40) that are representative of SRA datasets. Then, from each GTEx tissue dataset with at least 70 samples, we randomly selected samples to create ten datasets for each sample size (see *Methods*). We then applied all 30 workflows to construct coexpression networks from each one of these datasets. The resulting 72,900 networks were used to investigate the effects of varying each experimental factor by counting the number of times a given workflow significantly outperformed any other workflow (**Fig. 5**). In addition to this analysis with these resampled data, we also examined the effect of sample similarity and number of mapped reads directly in the SRA data by splitting the datasets into five equal size bins based on each of these factors and determining the number of times a given workflow was significantly better than another within each bin (**Fig. S7**).

**Figure 5.**
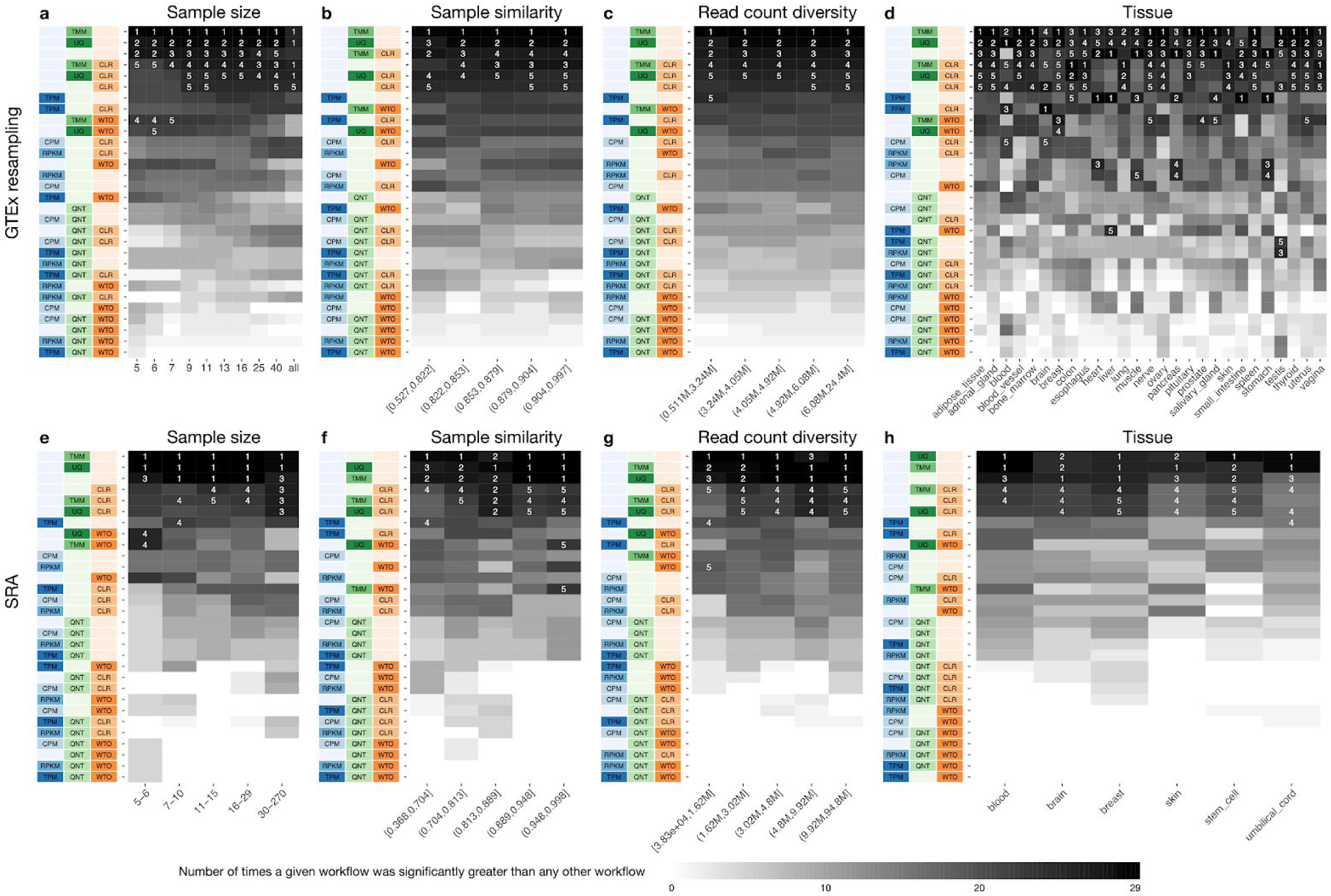
Impact of various dataset-related experimental factors on performance of workflows. Each heatmap shows the number of times (cell color) each workflow (row) outperforms other workflows as a particular experimental factor pertaining to the input datasets is varied (columns), when the resulting coexpression networks are evaluated based on the tissue-naive gold standard. The darkest colors indicate workflows that are significantly better than the most other workflows. In addition, the top 5 workflows in each column are marked with their rank, with ties given minimum rank. The heatmaps on the top (**a–d**) correspond to datasets from GTEx resampling and those on the bottom (**e–h**) correspond to SRA datasets. The heatmaps from left to right show workflow performance by sample size (**a, e**; number of samples used to make the coexpression network), sample similarity (**b, f**; median spearman correlation of 50% most variable genes between samples), read-count diversity by counts (**c, f**; standard deviation of counts sums across samples), and tissue of origin (**d, h**). *Figure S7* contains these heatmaps based on the tissue-specific gold standard.

In the GTEx-resampled data, *TMM* was significantly better than all other workflows for sample sizes 5 through 40 when using the naive standard for assessment (**Fig. 5**). *UQ* is a close second, performing significantly better than all workflows other than *TMM* at sample sizes 7 through 40. Using only *Counts* (no normalization) is surprisingly effective, especially at lower sample sizes, while *TMM_CLR* and *UQ_CLR* improve performance with increasing sample size. In fact, when all samples from a given GTEx tissue are used (≥70 samples), there is no significant difference between *TMM, UQ, TMM_CLR*, and *UQ_CLR. CLR* is the next best workflow after those top four. The only other workflows that are ever ranked in the top five are *TMM_WTO* and *UQ_WTO*, and that too only at low sample sizes (5–7). Based on the tissue-specific standards, *TMM_CLR* is the most effective workflow on all sample sizes except 5, where *TMM* and *UQ* are the top workflows (**Fig. S7**). For the highest two sample sizes (25 and 40), *TMM_CLR* is substantially better than all other workflows. The only workflows ranked in the top five in sample sizes 5 through 40 are *TMM_CLR, UQ_CLR, CLR, UQ* and *TMM. TMM* and *UQ* also perform well on the SRA data evaluated on the naive standard, being the top workflows in all five sample size groups (**Fig. 5**). Performance on the tissue-specific standards is more variable, with *Counts, TMM*, and *UQ* being top ranked in lower sample size groups and *CLR, UQ_CLR*, and *TMM_CLR* performing better in high sample size groups (**Fig. S7**). Again, it is clear that TMM and UQ are superior methods, with CLR improving performance in select cases.

Sample similarity and read-count diversity analyses show similar results to those from sample size analysis. When evaluating the GTEx-resampled data on the naive standard, *TMM* is almost always significantly better than every other workflow across all groups, while evaluating on the tissue-specific gold standards ranks *TMM_CLR* as the top workflow most consistently (**Fig. 5, Fig. S7**). In both standards, *TMM, UQ, CLR, TMM_CLR, UQ_CLR* and *Counts* are the workflows consistently showing up in the top five ranks. The SRA networks evaluated on either standard have *TMM, UQ*, and *Counts* showing up in the top three ranks across most groups, with *CLR, TMM_CLR*, and *UQ_CLR* making up most of the other workflows in the top five ranks (**Fig. 5, Fig. S7**).

Tissue is the factor that shows the most variability in terms of what makes up the top workflows, especially when evaluating on tissue-specific gold standards. This is due in part to the fact that splitting experiments by tissue results in the smallest groups, making significance more difficult to detect. Nevertheless, the top workflows from the analyses of other factors still have the best overall performance across all tissues. In the GTEx-resampled data, *TMM* is the top-ranked workflow most often based on the naive gold standard. *UQ* and *Counts* are almost always in the top five most significant workflows, while *TMM_CLR, UQ_CLR*, and *CLR* show up often. When evaluated on tissue-specific gold standards, *TMM_CLR* is ranked number one more frequently than any other workflow, but is not as consistent as *TMM* in the naive standard. *UQ_CLR, CLR, TMM*, and *UQ* are the other top-performing workflows, but a handful of other workflows enter the top five ranks in at least a few tissues. For SRA, only tissues that had more than fifteen separate experiments were used in the significance analysis (**Fig. S1**). On the naive standard, *UQ, TMM*, or *Counts* were always the most significant workflow in any given tissue and *CLR, TMM_CLR*, and *UQ_CLR* were usually in the top five. A similar pattern can be observed from the tissue-specific evaluations with the exception that *CLR* is ranked number one a few times. Taken together, these results suggest that the top-performing methods are largely robust to common experimental factors that vary from experiment to experiment. This property is critical because, to be practically beneficial, the best workflow for constructing coexpression networks should result in accurate coexpression networks irrespective of variations in these experimental factors.

The general trends presented above are all based on network accuracy measured using a metric based on the area under the precision-recall curve (log2(auPRC/prior)). These trends also hold when network accuracy is measured using precision at low recall, which focuses on maximizing the number of functional gene pairs among the high-scoring gene pairs instead of focusing on recovering all functional gene pairs. Put another way, these trends described above hold even when a threshold is applied to the coexpression network to retain just the high-scoring gene pairs for subsequent analysis. For the sake of completion, we have also evaluated all networks using the area under the ROC curve (auROC). All these results based on three different evaluation metrics (log2(auPRC/prior), precision at 20% recall, and auROC) are available as a consolidated webpage at https://krishnanlab.github.io/norm_for_RNAseq_coexp that researchers can explore to easily examine the performance of various workflows based on the properties of their RNA-seq dataset.

## Discussion

Despite the utility and growing popularity of coexpression analysis of RNA-seq data, relatively little focus has been devoted to identifying the optimal data normalization and network transformation methods that result in accurate RNA-seq-based coexpression networks. Here, we present the most comprehensive analysis of the effects of commonly-used techniques for RNA-seq data normalizations and network transformation on gene coexpression network accuracy (**Fig. 1**). We implemented 30 network-building workflows – one for every combination of within-sample normalization, between-sample normalization, and network transformation methods – and we ran each workflow on hundreds of RNA-seq datasets from GTEx and SRA. The resulting coexpression networks were evaluated using both known tissue-naive and tissue-specific gene functional relationships to ensure that the networks were tested for capturing not just generic but also gene interactions relevant to the tissue of interest (**Fig. S8**). The evaluations shed light on several key aspects of the impact of within-sample normalization, between-sample normalization, and network transformation methods (and their interplay) on the accuracy of the resulting coexpression networks.

### Impact of within-sample normalization

Within-sample normalization – commonly executed by converting gene counts to CPM, RPKM, or TPM – corrects for factors such as library size and gene length. As gene length biases both gene counts and their downstream analysis [18], it is not very surprising that TPM usually outperforms CPM, as CPM only corrects for library size and not gene length. However, the order in which gene-length and library-size correction are combined appears to be important. For example, studies have shown that RPKM, which first corrects for library size and then for gene length, is inferior to other methods in differential expression analysis and is not recommended [6–8]. Some studies have also noted that using RPKM does not necessarily take away the length bias in gene expression and can be unduly influenced by relatively few transcripts [6,19]. TPM was proposed as an improvement over RPKM by first correcting for length and then by library size. Thus, the resulting expression values more accurately reflect the “relative molar concentration” of an RNA transcript in the sample [20]. TPM normalization scales every sample to the same total RNA abundance (i.e. the same total sum of TPM values). Thus, gene expression across samples becomes more comparable when TPM normalized than when RPKM normalized. Consistent with these previous studies, we find that RPKM generally results in lower-performing coexpression networks and that TPM consistently outperforms CPM and RPKM, and can even occasionally perform better than the general top-performers TMM and UQ. Finally, since a number of technical and biological factors affect the size and makeup of the sample library, TPM has been found to be most effective when comparing samples from the same tissue type and experiment [21]. This observation could explain the good performance of TPM in our work wherein only samples within a dataset are compared and analyzed together to construct a coexpression network.

### Impact of between-sample normalization

Next, our results reinforce the expectation that between-sample normalization (using techniques such as TMM and UQ) leads to the largest improvement in coexpression accuracy. These methods are designed to make expression values across samples more comparable to one another, an aspect critical for coexpression analysis. However, QNT, a between-sample normalization method that is most commonly used with microarray data, performs very poorly for RNA-seq data. This is likely because QNT forces the distribution of samples to be exactly the same, meaning that each gene value is forced to be a particular quantile value. Consequently, it does not suit situations where there truly are different numbers of genes that are expressed outside of the typical ranges across samples [22,23], an effect that is further exacerbated in RNA-seq data because it has a larger dynamic range than microarray data. Genes with extreme values would not influence TMM or UQ normalization because they are explicitly excluded from the calculation. TMM specifically finds a subset of genes that are probably not differentially expressed between samples to make gene values comparable across the entire group, while UQ uses only the upper quartile gene values to adjust samples. This makes both normalizations robust to a number of highly or lowly expressed genes. However, large-scale changes in gene expression or high amounts of asymmetry, e.g. a large difference in the number of genes expressed above the typical range versus expressed below the typical range, violate these assumptions [22]. In our test cases, TMM and UQ performed the best, but it is possible that violation of their base assumptions may occur in specific disease conditions or external perturbations, leading to a significant decrease in their performances.

### Impact of network transformation

Network transformation is where there is most disagreement between GTEx and SRA data. CLR was the best network transformation method for GTEx data, while doing no transformation of the coexpression values gave the best results for SRA data. The most pronounced factor that explains this difference is sample size. The median sample size of SRA datasets is 12, while that of GTEx datasets is 197. Only four GTEx datasets have less than 70 samples (**Fig. S1**). Furthermore, GTEx resampling analysis showed that *TMM_CLR* and *UQ_CLR* improve with increasing sample size on the naive standard (**Fig. 5**) and to a lesser extent on the tissue-specific standards (**Fig. S7**) since CLR tended to already have better performance in general on tissue-specific standards than on the naive gold standard. For each gene pair, CLR adjusts the edge weight based on its value in relation to the distribution of edge weights for the individual genes in that pair to all other genes in the network. So, our hypothesis is that having a larger sample size results in a better estimate of each edge weight as well as the distribution of edge weights for each gene, which in turn increases CLR’s accuracy. Supporting this hypothesis, other studies have noted an association between larger sample size and more accurate coexpression networks [11,19], and subsequent network transformation with CLR [24]. WTO, on the other hand, performs poorly for both GTEx and SRA data. WTO adjusts the edge weight between gene pairs based on whether they share strong connections to the same set of genes in the network. Therefore, while CLR relies on summary statistics (mean and standard deviation) of edge distributions to adjust the edge weight between each gene pair, WTO relies on the actual, likely noisy, coexpression values, which may contribute to its inferior performance. It is also possible that CLR’s strategy more effectively deals with the mean-correlation relationship bias, or the observation that highly expressed genes tend to be more highly coexpressed, by capturing them as summary statistics, without relying on the fact that each of the correlation estimates are correct [25,26]. This may, in turn, explain why CLR tends to perform better on tissue-specific gold standards than on our naive gold standards, since genes that are ubiquitously expressed (and therefore involved in general, tissue-naive interactions) tend to be more highly expressed [27].

### Recommendations for building coexpression networks from RNA-seq data

By constructing coexpression networks for diverse datasets from both GTEx and SRA, we were able to evaluate workflows on large, homogeneous datasets as well as smaller, heterogeneous datasets to identify methods that are robust to differing technical and biological factors. Although there is some variation in performance between GTEx and SRA data, and slightly more variation introduced by tissue-specific gold standards, many trends are consistent across datasets and evaluations. Based on all our results, we make the following recommendations for building coexpression networks from RNA-seq data:

- If gene counts are available, use TMM or UQ to normalize the data. They consistently give the best performance regardless of various factors. Between the two, TMM seems to be slightly more consistent in yielding top performance. Even though no normalization (Counts) leads to good performance in our study, applying the additional normalization step is prudent to ensure robustness against variabilities specific to a new dataset.
- If data is only available after within-sample normalization, use TPM for coexpression analysis. Data in CPM and RPKM units can be easily converted to TPM. TPM outperforms CPM and RPKM and yields consistently reasonable performance.
- If the dataset has greater than 40 samples, use CLR to transform the pairwise gene correlations. CLR may also help certain cases where the main interest is interactions that are tissue-specific.
- QNT and WTO hurt performance in combination with every other method, in all cases, and should not be used.

To enable researchers to explore all the underlying results in a streamlined manner and find the analyses that are relevant to coexpression analysis of their own RNA-seq datasets, we have made them available as a rich webpage written with R Markdown:

https://krishnanlab.github.io/norm_for_RNAseq_coexp.

### Potential future applications and extensions

Going forward, we can leverage this comprehensive benchmarking framework for coexpression analysis to answer newer and subtler questions about data quality and sample composition. For example, many studies have found that removing unwanted variation, i.e. noise caused by technical rather than biological factors, in the RNA-seq data has led to improvements in downstream analysis including the calculation of coexpression networks [28,29]. Such corrections are often done using SVD-based methods, including removing the first (or the first few) principal components. However, caution must be taken when using these methods as they may easily remove biological signals from the data [30], especially in typical small-to-medium-sized datasets produced by most research labs (e.g. represented in SRA). Future work using our framework could help learn the guidelines for deciding which and how many factors to remove while carefully considering the various properties of the data and the biological objective of the analysis. For instance, one could explore if different tissues might be sensitive to different technical factors; signal from blood is often heavily influenced by the large variation in cell type composition but the brain is much more greatly affected by the post-mortem-interval [31]. Another related and open question is how cell type composition influences gene coexpression calculated from bulk tissue data. Some studies have concluded that gene coexpression networks are heavily confounded by this factor [32,33], while others have shown that coexpression derived from single-cell data is very similar to bulk coexpression [34,35]. Finally, a similar framework could also be used to explore the best procedure for building coexpression networks from single-cell RNA-seq data, which has an entirely different set of challenges [36] that call for an entirely separate benchmarking effort.

## Conclusions

We have performed an extensive benchmarking and analysis of how data normalization and network transformation impact the accuracy of coexpression networks built from RNA-seq datasets. Based on this work, we have arrived at concrete recommendations on robust procedures that will generally lead to best coexpression networks. Specifically, using trimmed mean of M-values (TMM) and upper quartile (UQ) normalizations to construct coexpression networks results in the most consistently high accuracy networks, and using CLR to transform the network can further increase accuracy in select cases. All the results from this study – for GTEx, SRA, and GTEx resampling datasets, based on tissue-naive and tissue-specific gold standard, using three different evaluation metrics – are available as a consolidated webpage at https://krishnanlab.github.io/norm_for_RNAseq_coexp. Researchers can use this website to easily examine the performance of various workflows and make appropriate choices for coexpression analysis based on the properties of their RNA-seq dataset of interest.

## Methods

### Data Collection

Read counts for both SRA and GTEx datasets were downloaded from the Recount2 database [12] and processed separately. Recount2 aligns all sequenced reads using Rail-RNA, which eliminates the effect of using different alignment software on separate experiments. We obtained SRA data for any tissue with at least five separate experiments that each had at least five samples. The set of samples from each experiment (project) was considered as an individual dataset from which coexpression networks are inferred (one network per dataset). If a given experiment had samples from multiple tissues, the samples were divided into multiple datasets that each contain samples from the same tissue to yield 543 candidate SRA datasets. We downloaded all available GTEx data, which was a total of 9,657 samples from 31 tissues.

### Preprocessing

As a form of quality control, we excluded experiments that Recount2 identified as having a misreported paired-end status. Experiments that contained “cell line”, “celll line”, “passage”, “cultured cells”, or “cell culture” in the characteristics metadata were also removed so as to retain primary tissue samples, which left 341 SRA datasets. Next, we discarded low-coverage samples that had zero expression (counts) in at least half of all genes of interest (lncRNA, antisense RNA, and protein-coding genes), and subsequently excluded entire datasets that no longer contained five or more samples. Retaining only tissues that still had at least 5 separate experiments left 256 datasets. Finally, we removed genes with very low expression across the board by filtering out those that did not have at least one read per million sample reads in at least 20% of the samples in at least one dataset. This resulted in 22,084 genes in the SRA networks and 20,418 genes in the GTEx networks.

### Calculating gene counts

Recount2 stores quantified expression as base pair counts per gene. We converted these values into gene counts by dividing these base pair per gene counts by the average read length in the sample and accounted for paired-end read samples by further dividing by a factor of two.

### Within-sample normalization

Within-sample normalization is designed to transform the expression levels of genes within the same sample so that they can be compared to each other. Here, we considered counts per million (CPM), transcripts per million (TPM), and reads per kilobase million (RPKM) for performing within-sample normalization of the original raw gene counts [20,37]. Note that RPKM is almost the same as Fragments Per Kilobase Million (FPKM), except FPKM was introduced to accommodate paired-end RNA-seq so it accounts for the fact that two reads can map back to a single fragment. We account for paired-end samples with FPKM, but use the term “RPKM” throughout the manuscript. These three ways of normalizing counts are very commonly used in RNA-seq analysis and account for library size and gene/transcript length in different ways. CPM corrects for library size (expressed in million counts) so that each count is expressed as a proportion of the total number reads in the sample. TPM and RPKM are similar methods that correct for both library size and gene length. Each gene count is divided by both the length of the gene and the sum of counts in the sample, but these operations are done in a different order. TPM divides counts by gene length (in kb) first to get transcript counts and then by total number of transcripts in the sample, resulting in each normalized sample having the same number of total counts. This is not guaranteed for RPKM since it corrects each gene count for the total number of reads in the sample before correcting for gene length.

### Between-sample normalization

Between-sample normalization transforms the expression levels of genes across a group of samples so that gene counts from the same gene in different samples can be more accurately compared to each other. We tested quantile (QNT), trimmed mean of M-values (TMM) [38], and upper quartile (UQ) normalizations [6]. Quantile normalization is an extremely popular between-sample normalization for microarray samples. Applied to RNA-seq data, QNT forces the distribution of all gene expression values to be exactly the same in each sample. We performed quantile normalization on counts, CPM, TPM, and RPKM using the *preprocessCore* package available from Bioconductor. TMM normalizes across samples by finding a subset of genes whose variation is mostly due to technical rather than biological factors, i.e. not differentially expressed, then using this subset to calculate a scaling factor to adjust each sample. In brief, each sample is compared to a chosen reference sample. A certain upper and lower percentage of data based on absolute intensity and log-fold-change relative to the reference sample is removed (by default, 5% for absolute intensity and 30% for log-fold-change) and the log-fold-changes of the remaining set of genes are used to calculate a single scaling factor for the non-reference samples. UQ normalization first removes all zero-count genes and calculates a scaling factor for each sample to match the 75% quantile of the counts in all the samples. In both TMM and UQ, the scaling factors are adjusted to multiply to one before they are used to normalize each sample. We performed trimmed mean of M-values (TMM) and UQ normalization of the counts data using the *edgeR* package.

### Network construction

A coexpression network was constructed for each individual dataset by calculating the Pearson correlation coefficient between every pair of genes in that dataset using the *Distancer* tool in the *Sleipnir* C++ library. These correlations were treated as the edge weight between gene pairs. We chose Pearson correlation as it has been repeatedly shown to provide a robust measure of gene-gene correlations, especially in small-to-medium-sized datasets that are produced by individual laboratories [39,40].

### Network transformation

Coexpression networks are noisy and indiscriminately capture indirect interactions due to being calculated from noisy, steady-state gene expression data. Hence, several studies have proposed methods to modify the raw coexpression network to upweight connections that are more likely to be real and downweight spurious correlations based on the topology of the network. As correctly normalized RNA-seq data should yield more accurate quantification of gene activities, successfully transformed networks should yield more accurate estimates of functional relationships between genes. We tested two common methods of network transformation, weighted topological overlap (WTO) [41] and context likelihood of relatedness (CLR) [42], that use different aspects of network topology to correct the raw coexpression network. The general idea of WTO is to increase the edge weight between gene pairs that share a high number of network neighbors while diminishing edge weight between gene pairs that are tightly connected to very different sets of genes in the network. All edges in the resulting network will have normalized weighted between zero and one. CLR reweights the edge for each gene pair (*i, j)* based on how different the original weight of that edge is relative to all of the connections to gene *i* and all connections to gene *j* (to the rest of the genes in the network). For instance, CLR will upweight an edge between two genes if the edge weight is high compared to all of the other connections of both genes. WTO was implemented using the *wTO* function with the “sign” method in the *wTO* package [43] and CLR was implemented using the *Dat2Dab* function in the *Sleipnir* C++ library.

### Network evaluation

The goal of coexpression networks is to capture true functional relationships between genes in the cellular context of the original dataset. Therefore, we evaluated the accuracy of each coexpression network by comparing it to two gold standards, one representing known generic (tissue-naive) functional relationships and the other representing known tissue-specific gene functional relationships. We assembled these gold standards by beginning with a set of manually-selected Gene Ontology Biological Process (GOBP) terms [39] that were deemed to be specific enough to be confident that any genes co-annotated to them could be considered functionally related via experimental follow-up. Then, any pair of genes that were co-annotated to the same specific GOBP term was set as a positive edge in the gold standard. We only used annotations based on experimental (GO evidence codes: EXP, IDA, IPI, IMP, IGI, TAS) or curated evidence (IC). We explicitly ignored gene-term annotations made based on expression (GO evidence code: IEP) to avoid circularity when comparing coexpression-derived interactions to this gold-standard. We next had to determine which pairs of genes among the ones with at least one positive edge could be declared as negative edges. Following previous work, we ignored gene pairs not co-annotated to any specific term but still interact with many of the same genes in the gold standard (determined based on each being annotated to two different terms that contained very similar sets of genes; hypergeometric test; p-value <0.05). We also ignored gene pairs that were not co-annotated to any specific term but were co-annotated to certain general GOBP terms, thus introducing ambiguity in whether they are functionally related or not. All remaining gene pairs were considered negatives. We built the naive gold standard using the *Answerer* function in the *Sleipnir* C++ library.

We created the tissue-specific gold standards for as many tissues as possible by subsetting the naive gold standard based on genes known to be specifically expressed in a particular tissue. We obtained tissue-specific genes from the TISSUES 2.0 database, including those that had a z-score ≥ 3.5 in the Knowledge channel [44]. For a given tissue, a positive edge from the naive gold standard was kept in its tissue-specific standard if both genes were expressed in that tissue. Negative edges were kept if both genes were expressed in that tissue, or if one gene is expressed in the tissue and the other gene is expressed in one of the other tissues considered. Only standards containing at least 50 positive edges were used for evaluation, resulting in 24 tissue-specific gold-standards. We specifically excluded epithelium from consideration for a tissue-specific standard, as there is no straightforward way to determine the body site each sample was taken from.

We used the *DChecker* function in the *Sleipnir* C++ library to compare each coexpression network to each gold-standard and return the number of true positives, false positives, true negatives, and false negatives at various edge weight thresholds. These numbers were used to calculate the area under the precision-recall curve (auPRC) using the *trapz* function in the *pracma* package. Since gene functional relationship gold-standards of different tissues have different proportions of positives to negatives, the original auPRC scores are not directly comparable to each other. Therefore, we divided each auPRC by its “prior” – the auPRC of a random predictor, equal to the fraction of positives among all positive and negative edges – and expressed the performance as the logarithm of this ratio to enable tissue-to-tissue comparisons.

### Workflow comparison and analysis by parts

To assess whether two workflows resulted in coexpression networks that were significantly different in quality, we used a paired Wilcoxon rank sum test to compare the auPRC scores across all coexpression networks generated by those two workflows. After calculating p-values, we performed a correction for multiple testing with the Benjamini-Hochberg procedure and declared workflows with FDR ≤ 0.01 as being significantly different. Further, each workflow is a combination of method choices at multiple stages. So, to determine the impact of including a particular method in a workflow, we across aggregated workflows to calculate the proportion of times that including a particular method in a workflow resulted in the workflow being significantly greater than one that did not include the method. As it is not possible to do within-sample normalization and then do either TMM or UQ, any workflow including CPM, TPM, or RPKM was excluded when assessing between-sample normalization methods so that method being compared to each other based on the same number of aggregated workflows. For similar reasons, workflows involving TMM and UQ were not considered for the analysis of within-sample normalization methods.

### GTEx resampling

To simulate uniformly-processed datasets that have sample sizes similar to datasets from SRA, we chose nine sample sizes (5, 6, 7, 9, 11, 13, 16, 25, and 40) based on the distribution of SRA dataset sample sizes. Then, from each GTEx dataset with at least 70 samples, we randomly sampled a “dataset” of each sample size, repeating this sampling ten times to create 10 datasets per sample size from each GTEx dataset. One coexpression network was constructed and evaluated from each of these GTEX-resampled datasets in the same manner outlined above.

### Experimental factor analysis

In addition to dataset size (i.e. number of samples), the quality of the coexpression network reconstructed from a dataset could also depend on the similarity between the samples in that dataset as well as the total number of mapped reads. We performed an analysis to determine this impact using the GTEx-resampled datasets and the original SRA datasets. Since SRA datasets are not large enough to do resampling for sample size analysis, we split them into five groups with equal number of datasets, with datasets in each group having similar sample sizes. We define sample similarity for a given dataset as the median spearman correlation between all samples using the 50% most variable genes in the GTEx tissue they came from for the resampled GTEx datasets, or the median spearman correlation between all samples using the 50% most variable genes in each individual dataset in the case of the SRA networks. Read-count diversity is calculated by summing the gene counts of each sample in a given dataset and taking their standard deviation. Based on each of these measures – sample similarity and read-count diversity – we divided the datasets into five groups of equal size while taking care to check that each group contained datasets with similar sample sizes. For the tissue analysis, we could only determine significance in SRA tissues that had at least 15 datasets.

## Declarations

### Ethics approval and consent to participate

Not applicable

### Consent for publication

Not applicable

### Availability of data and materials

The expression datasets used in this study can be obtained from the Recount2 database

https://jhubiostatistics.shinyapps.io/recount/.

All the results from this study are available at

https://krishnanlab.github.io/norm_for_RNAseq_coexp.

### Authors’ contributions

KAJ and AK designed the study. KAJ performed all the analyses. KAJ and AK interpreted the results and wrote the final manuscript.

## Acknowledgements

We thank the members of the Krishnan Lab for helpful discussion. We are particularly grateful to Anna Yannakopoulos and Chris Mancuso for code advice, and Stephanie Hickey for suggestions on the manuscript.

