## Supplemental Material for "Robust normalization and transformation techniques for constructing gene coexpression networks from RNA-seq data"

### Supplemental Figures

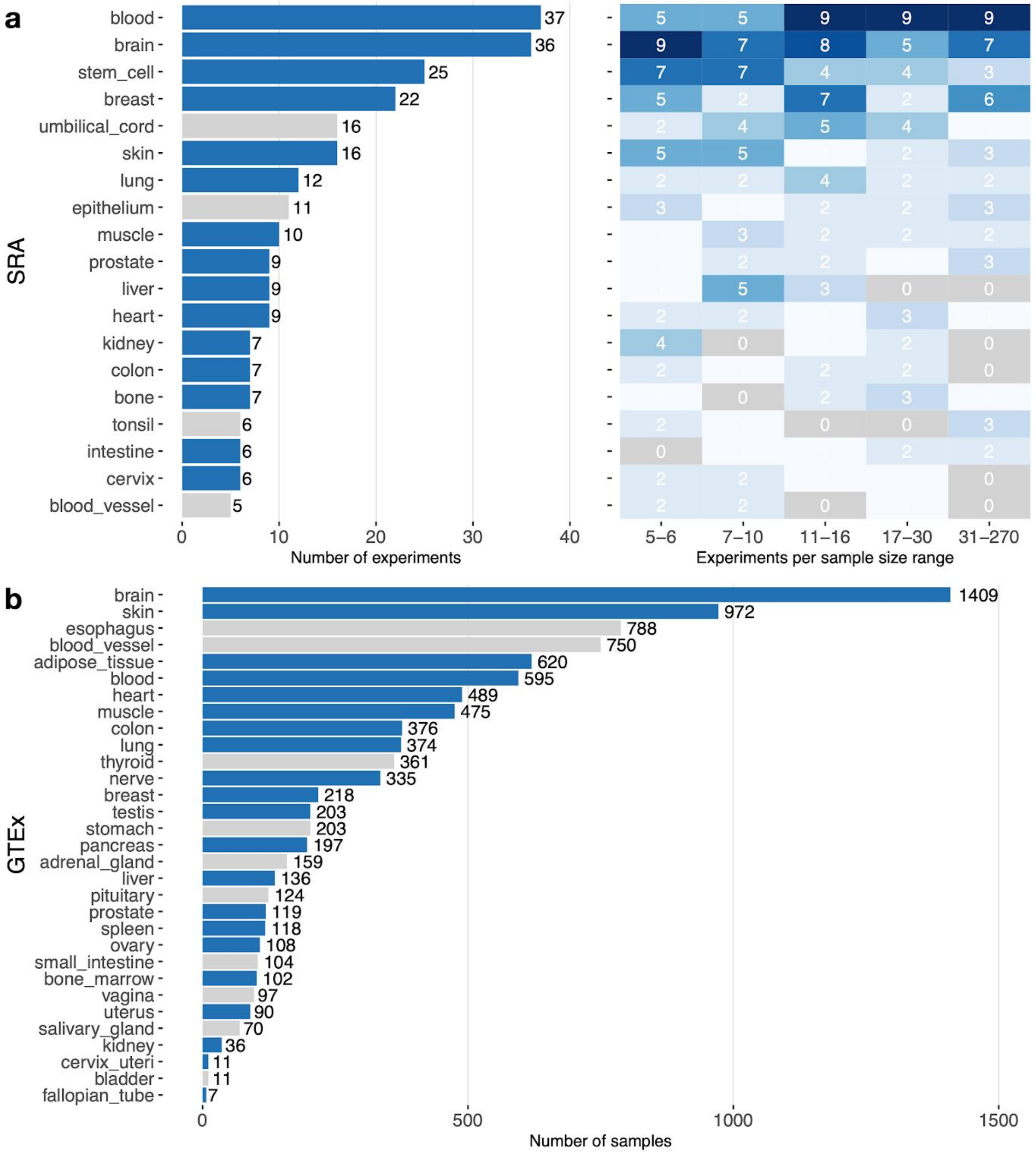


**Figure S1. Data used in this study.** (**a**) The barplot shows the number of experiments from each tissue in the SRA data. The heatmap on the right shows the number of projects/experiments that have a particular sample size for each tissue. (**b**) The barplot shows the number of samples for each GTEx tissue. In the barplots, blue bars indicate tissues for which we were able to create a tissue-specific gold standard. Tissues with gray bars were evaluated on the tissue-naive standard only.

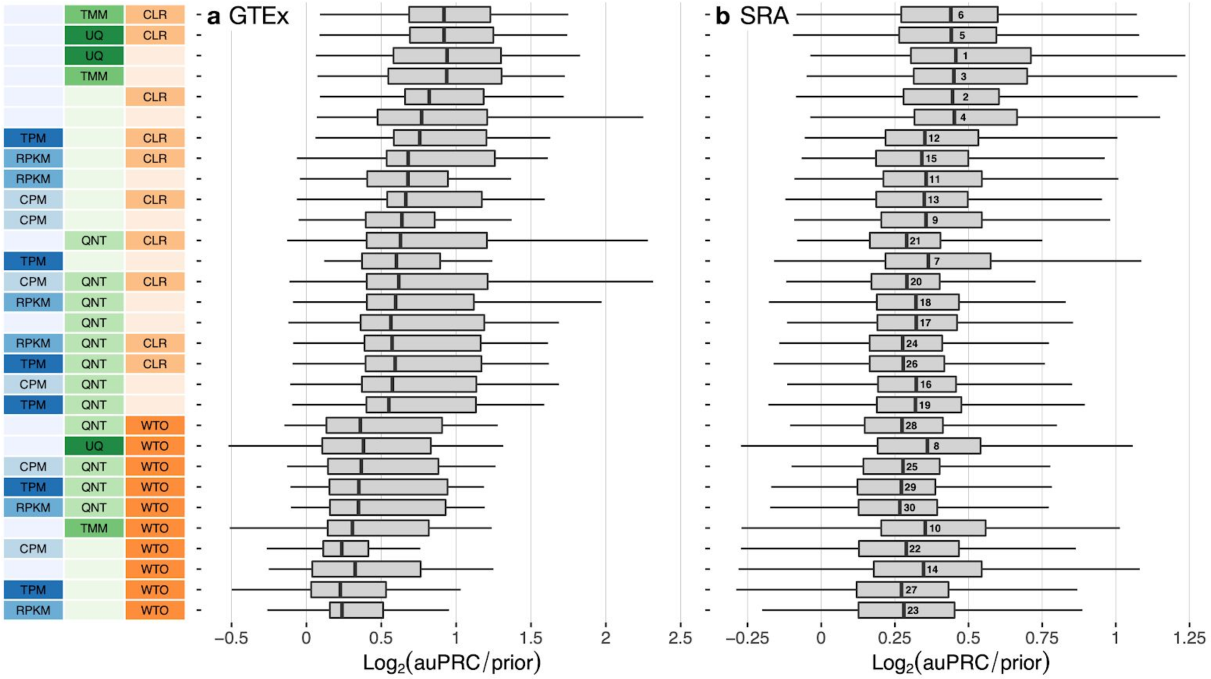


**Figure S2. Overall performance of workflows based on the tissue-specific gold standard.** The plots show the aggregate accuracy of all coexpression networks resulting from each individual workflow using (**a**) GTEx and (**b**) SRA datasets, evaluated based on the tissue-naive gold standard. The workflows (rows) are described in terms of the specific method used in the within-sample normalization (blues), between-sample normalization (greens), and network transformation (oranges) stages. The performance of each workflow is presented as boxplots that summarizes the log2(auPRC/prior) of each workflow where auPRC is the area under the precision recall curve (see Methods). The workflows are ordered by their median log2(auPRC/prior) for the GTEx data. The numbers inside the SRA boxes indicate rank by median log2(auPRC/prior) of the workflows for the SRA data. *Figure 2* contain these performance plots based on the tissue-specific gold standard.

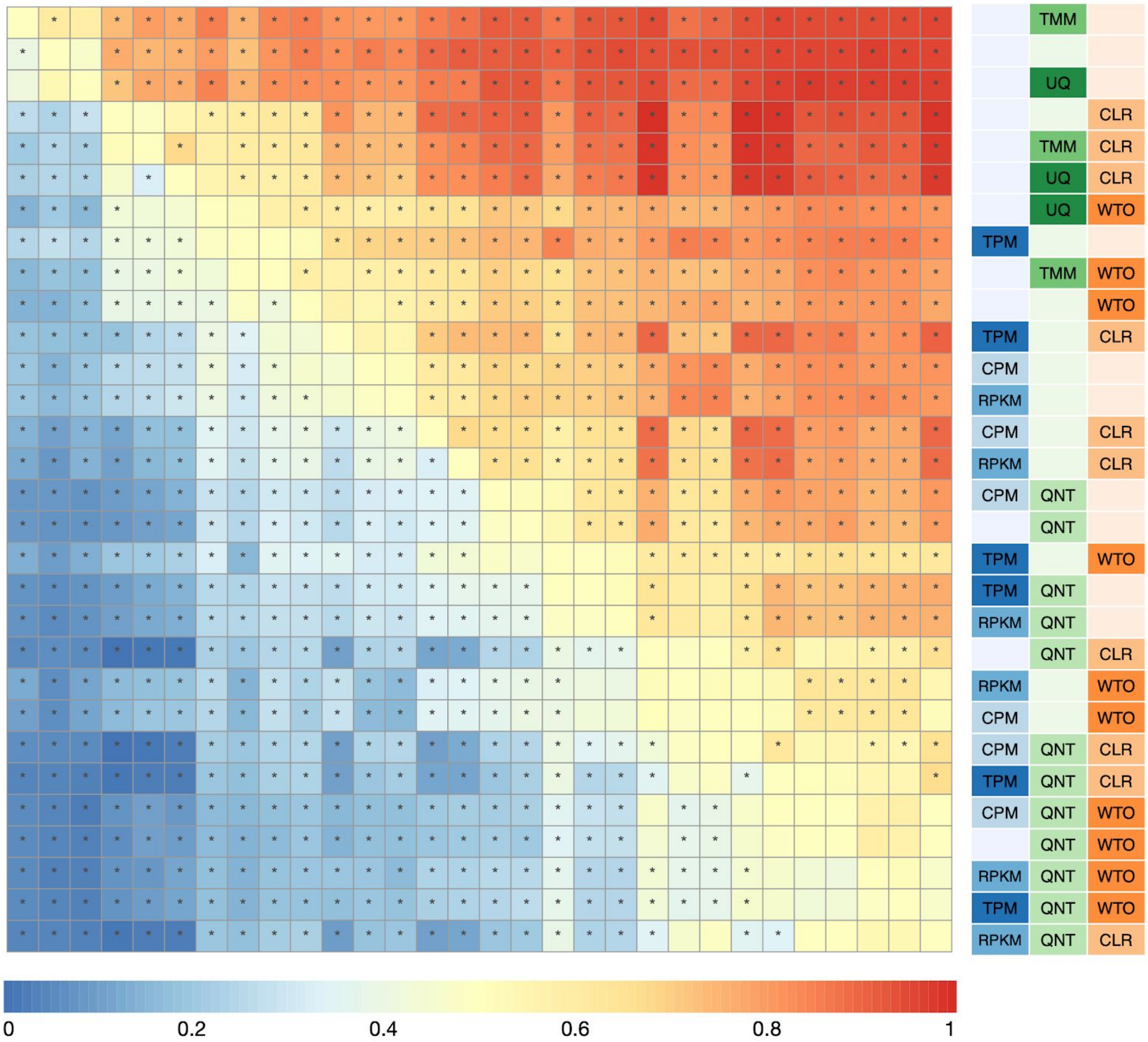


**Figure S3. Dataset-level pairwise comparison of workflow performance for SRA datasets based on the tissue-naive gold standard.** The heatmap shows the relative performance of a pair of workflows, corresponding to a row and a column, directly compared to each other for the SRA datasets based on the tissue-naive gold standard. The color in each cell (row, column) represents the proportion of datasets for which the workflow along the row has a higher log2(auPRC/prior) than the workflow along the column. Comparisons that are statistically significant (corrected $p < 0.01$) based on a paired Wilcoxon test are marked with an asterisk. *Figures 3a* contains the corresponding heatmap for GTEx datasets.

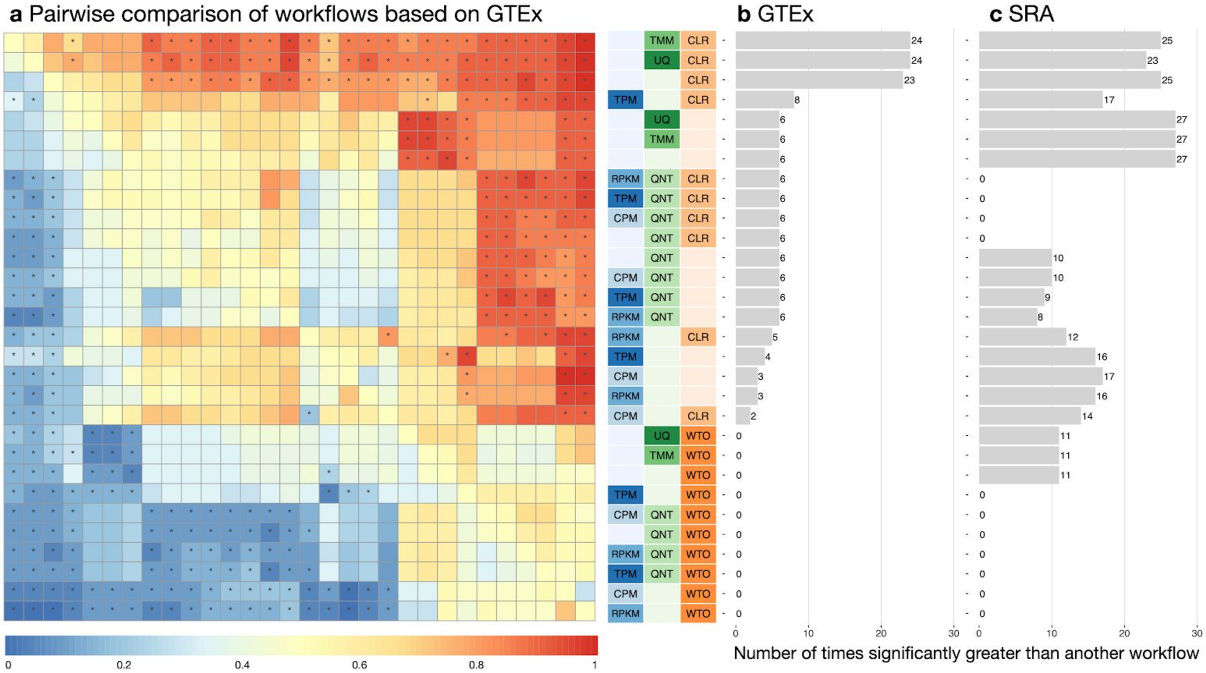


**Figure S4. Dataset-level pairwise comparison of workflow performance for GTEx and SRA datasets based on the tissue-specific gold standard.** (**a**) The heatmap shows the relative performance of a pair of workflows, corresponding to a row and a column, directly compared to each other for the GTEx datasets based on the tissue-specific gold standard. The color in each cell (row, column) represents the proportion of datasets for which the workflow along the row has a higher log2(auPRC/prior) than the workflow along the column. Comparisons that are statistically significant (corrected $p < 0.01$) based on a paired Wilcoxon test are marked with an asterisk. *Figures S4* contains the corresponding heatmap for the SRA datasets. (**b** and **c**) Barplots show the number of times each workflow was significantly greater than another workflow for GTEx (left) and SRA (right) datasets. Figure 3 and S3 contain these performance plots based on the tissue-naive gold standard.

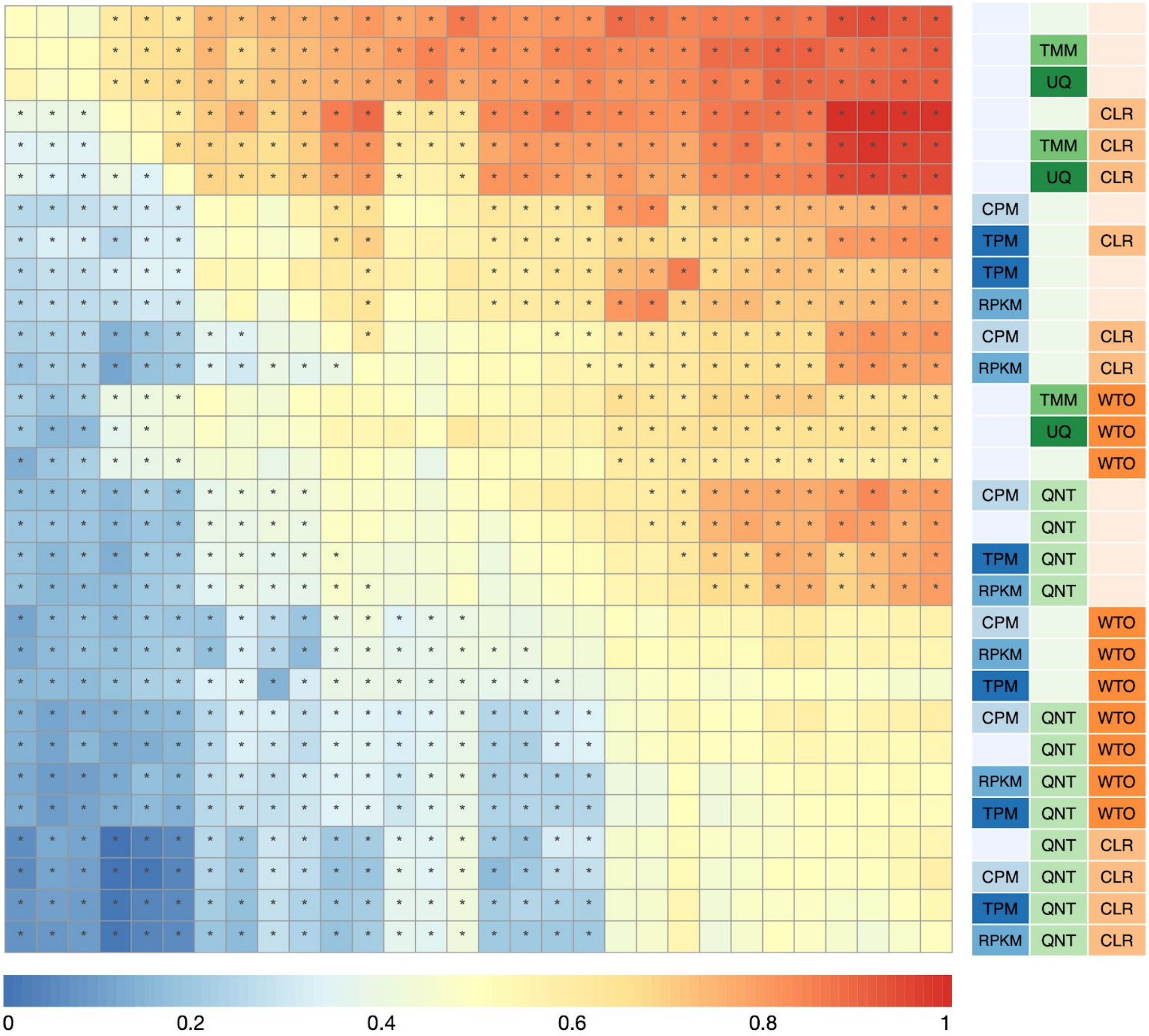


**Figure S5. Dataset-level pairwise comparison of workflow performance for SRA datasets based on the tissue-specific gold standard.** The heatmap shows the relative performance of a pair of workflows, corresponding to a row and a column, directly compared to each other for the SRA datasets based on the tissue-specific gold standard. The color in each cell (row, column) represents the proportion of datasets for which the workflow along the row has a higher log2(auPRC/prior) than the workflow along the column. Comparisons that are statistically significant (corrected $p < 0.01$) based on a paired Wilcoxon test are marked with an asterisk. *Figure S4a* contains the corresponding heatmap for GTEx datasets.

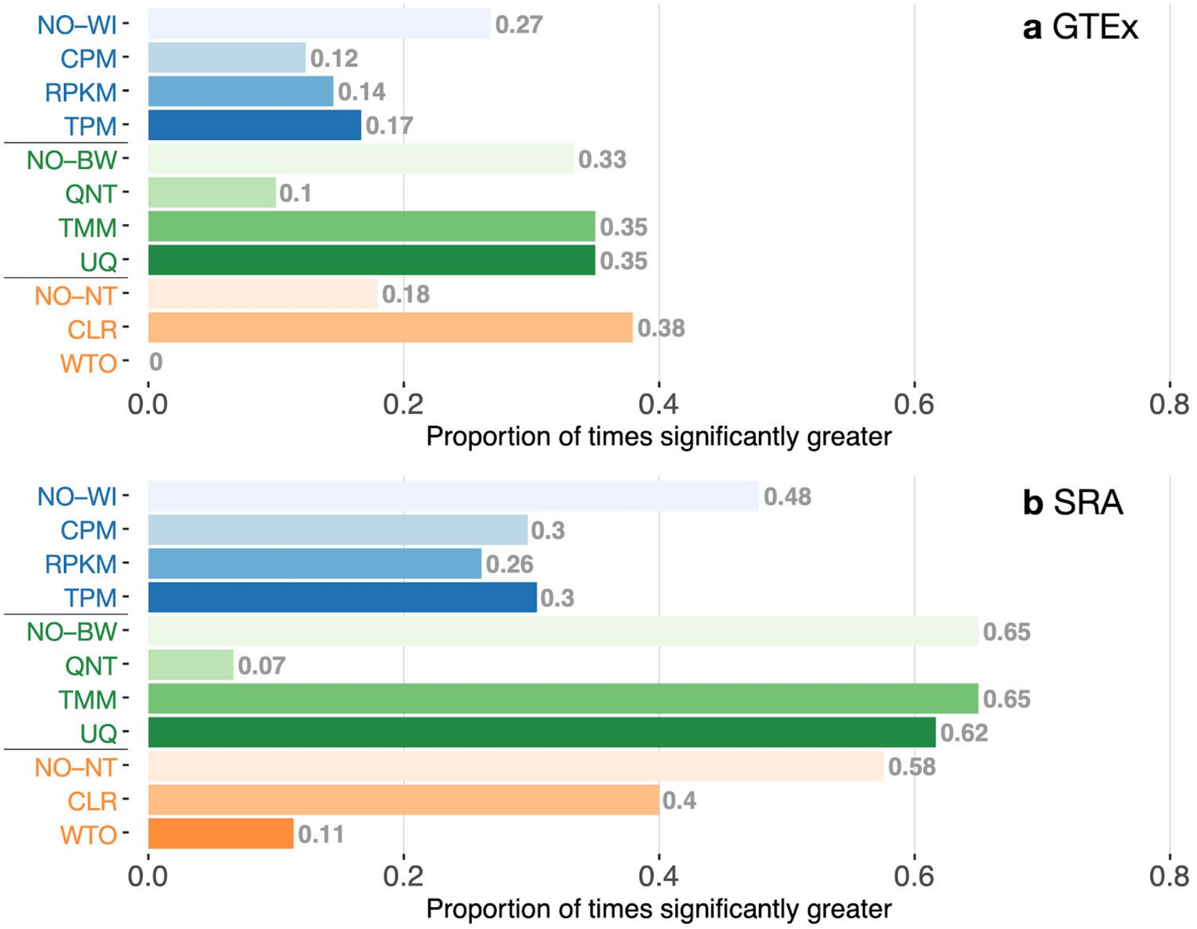


**Figure S6. Impact of individual methods on performance of workflows based on the tissue-specific gold standard.** Each bar in the two barplots, corresponding to a specific method, shows the proportion of times (x-axis) that workflows including that particular method (y-axis) were significantly better than other workflows. The barplots correspond to performance for the (**a**) GTEx and (**b**) SRA datasets evaluated on the tissue-naive gold standard. In order to make the comparison of between-sample normalization methods fair, workflows including CPM, RPKM, or TPM were left out because it is not possible to pair them with TMM or UQ normalization. Similarly, TMM and UQ methods are not included for “no within-sample normalization” (NO–WI). *Figure 4* contains these barplots based on the tissue-naive gold standard.

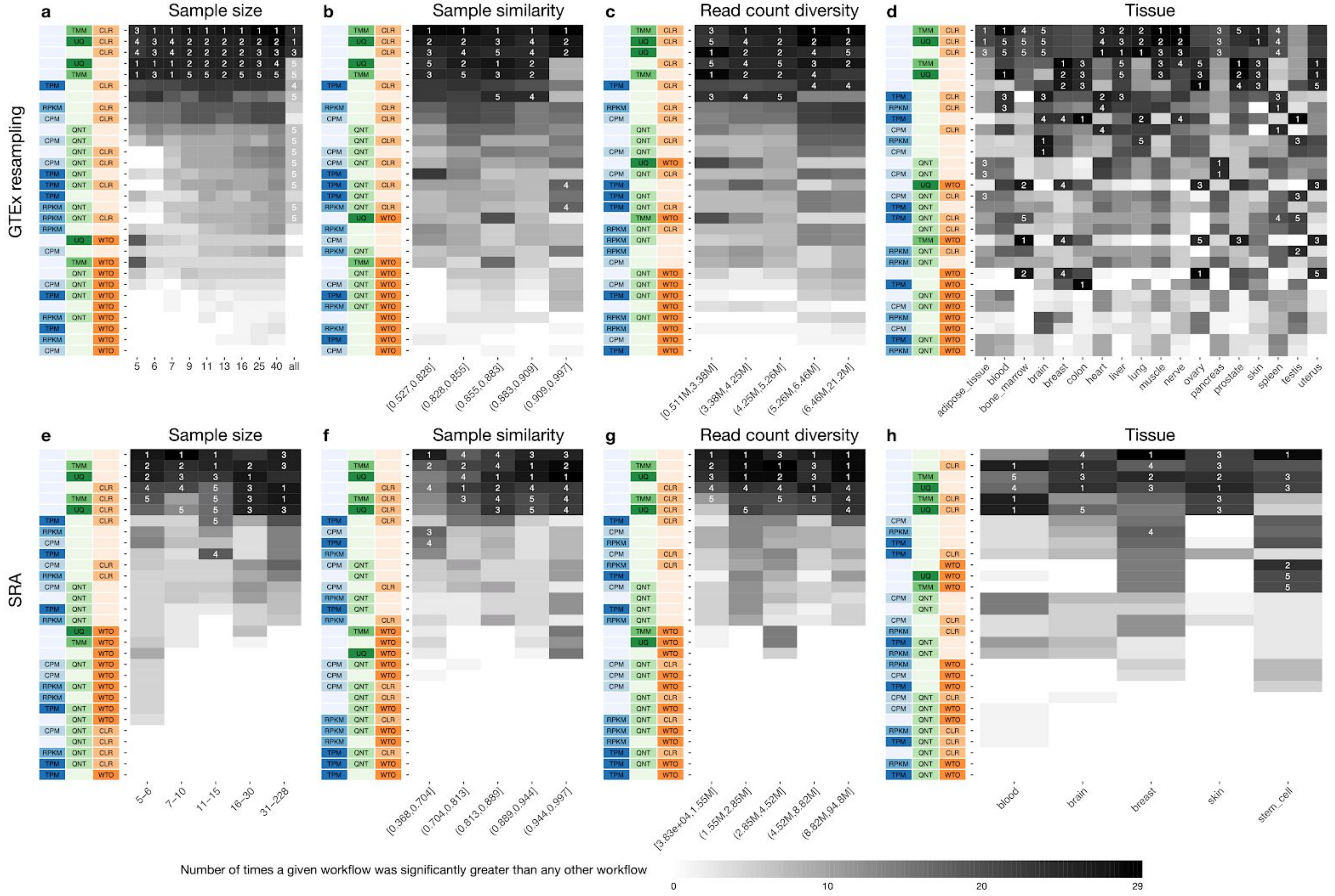
a
Sample size
b
Sample similarity
c
Read count diversity
d
Tissue
GTEx resampling
e
Sample size
f
Sample similarity
g
Read count diversity
h
Tissue
SRA
TMM
UQ
CLR
TPM
RPKM
CPM
QNT
WTO
5
6
7
9
11
13
16
25
40
all
(0.527,0.828]
(0.828,0.855]
(0.855,0.883]
(0.883,0.909]
(0.909,0.997]
[0.511M,3.38M]
(3.38M,4.25M]
(4.25M,5.26M]
(5.26M,6.46M]
(6.46M,21.2M]
adipose_tissue
blood
bone_marrow
brain
breast
colon
heart
liver
lung
muscle
nerve
ovary
pancreas
prostate
skin
spleen
testis
uterus
5–6
7–10
11–15
16–30
31–228
[0.368,0.704]
(0.704,0.813]
(0.813,0.889]
(0.889,0.944]
(0.944,0.997]
[3.83e+04,1.55M]
(1.55M,2.85M]
(2.85M,4.52M]
(4.52M,8.82M]
(8.82M,94.8M]
blood
brain
breast
skin
stem_cell
Number of times a given workflow was significantly greater than any other workflow
0
10
20
29

**Figure S7. Impact of various dataset-related experimental factors on performance of workflows based on the tissue-specific gold standard.** Each heatmap shows the number of times (cell color) each workflow (row) outperforms other workflows as a particular experimental factor pertaining to the input datasets is varied (columns), when the resulting coexpression networks are evaluated based on the tissue-naive gold standard. The darkest colors indicate workflows that are significantly better than the most other workflows. In addition, the top 5 workflows in each column are marked with their rank, with ties given minimum rank. The heatmaps on the top (**a**–**d**) correspond to datasets from GTEx resampling and those on the bottom (**e**–**h**) correspond to SRA datasets. The heatmaps from left to right show workflow performance by sample size (**a**, **e**; number of samples used to make the coexpression network), sample similarity (**b**, **f**; median spearman correlation of 50% most variable genes between samples), library size diversity by counts (**c**, **f**; standard deviation of counts sums across samples), and tissue of origin (**d**, **h**). *Figure 5* contains these heatmaps based on the tissue-naive gold standard.

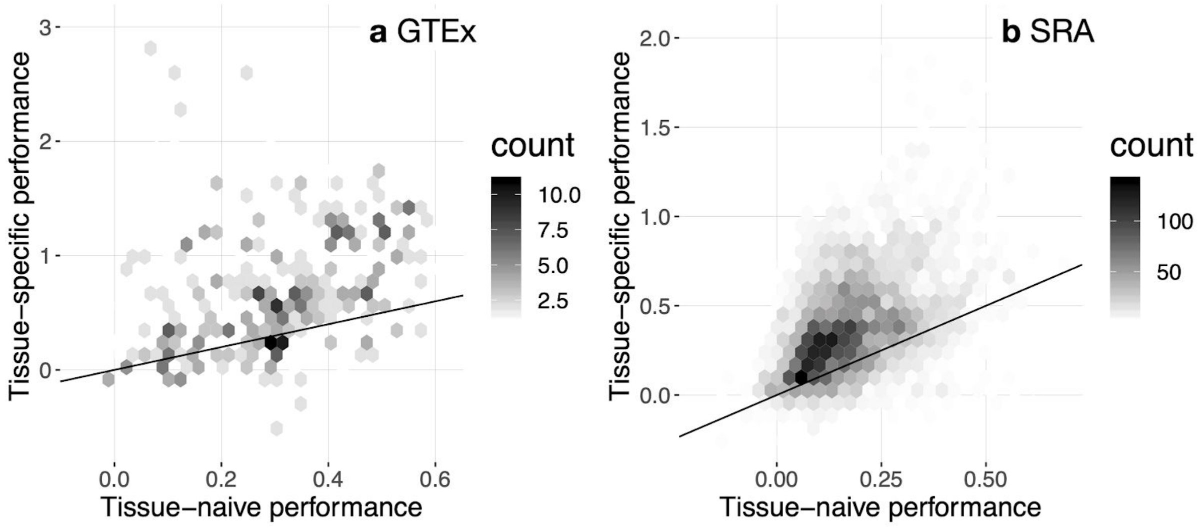


**Figure S8. Overall accuracy of coexpression networks when evaluated based on the tissue-naive and tissue-specific gold standards.** Each density plot – for the (**a**) GTEx and (**b**) SRA datasets – shows the distribution of log2(auPRC/prior) across all workflows and datasets when evaluating based on the tissue-naive gold standard (x-axis) vs. the tissue-specific gold standard (y-axis). These distributions show that coexpression networks capture tissue-specific gene interactions and emphasises the importance of evaluating coexpression networks using tissue-specific gold standards.
